# 25 Years of Pathway Figures

**DOI:** 10.1101/2020.05.29.124503

**Authors:** Kristina Hanspers, Anders Riutta, Martina Kutmon, Alexander R. Pico

## Abstract

**Background:** Pathway diagrams are fundamental tools for describing biological processes in all aspects of science, including training, generating hypotheses, describing new knowledge and ultimately as communication tools in published work. Thousands of pathway diagrams are published each year as figures in papers. But as static images the pathway knowledge represented in figures is not accessible to researchers for computational queries and analyses. In this study, we aimed to identify pathway figures published in the past 25 years, to characterize the human gene content in figures by optical character recognition, and to describe their utility as a resource for pathway knowledge.

**Approach:** To identify pathway figures representing 25 years of published research, we trained a machine learning service on manually-classified figures and applied it to 235,081 image query results from PubMed Central. Our previously described pipeline ^1^ was utilized to extract human genes from the pathway figure images. These figures were characterized in terms of their parent papers, human gene content and enriched disease terms. Diverse use cases were explored for this newly accessible pathway resource.

**Results:** We identified 64,643 pathway figures published between 1995 and 2019, depicting 1,112,551 instances of human genes (13,464 unique NCBI Genes) in various interactions and contexts. This represents more genes than found in the text of the same papers, as well as genes not found in any pathway database. We developed an interactive web tool to explore the results from the 65k set of figures, and used this tool to explore the history of scientific discovery of the Hippo Signaling pathway. We also defined a filtered set of 32k pathway figures useful for enrichment analysis.

## Introduction

The molecular mechanisms underlying biology are often outlined as pathway diagrams. In textbooks and on whiteboards, these depictions are fundamental to a biologist’s training. As mental models for practitioners, they serve as scaffolds for hypotheses and integrating new knowledge. And in the scientific literature, pathway figures are the pinnacle of communication for published work, synthesizing diverse sources and types of data spanning decades into a coherent model. Though often published only as static images, pathways express dynamic interactions. Common examples include metabolic cycles, gene regulation and signaling cascades. Depicted interactions play out over a spectrum of electrochemical, enzymatic and developmental timescales.

When properly modeled as an interaction network and annotated with standard identifiers, pathway knowledge can be conveyed with greater precision in formats amenable to computational analysis. Distinct from static images, pathway models can be used in enrichment analyses ^2^, enhanced data visualization ^3;4^, knowledge graphs ^5;6^, biomedical inference ^7^ and database queries ^8;9^. Over the past couple decades a number of pathway databases, including GenMAPP ^10^, MetaCyc ^11;12^, KEGG ^13^ and Reactome ^14;15^ took on the challenge of curating canonical pathway biology, each with their own unique focus and approach. A broader, community-curated approach was undertaken by WikiPathways to allow any researcher to model and freely share their pathway knowledge ^16;17;18^. And under an even broader umbrella, the NDEx database provides access to not only pathways, but also diverse types of network models, offering DOI minting for citation ^19^. Despite the continued growth and active usage of these database efforts, the vast majority of pathway knowledge is still captured in static images submitted solely to publishers as figures. We estimate 1,000 pathway figures are indexed by PubMed Central (PMC) *each month* in recent years and less than 3% of these are sourced from a pathway database ^1^.

In this study, we have identified pathway figures published over the past ^25^ years and characterized their content in terms of recognized gene symbols by optical character recognition (OCR). While it is more common to process text from the abstracts and body of papers in order to extract genes and other biological concepts including interactions, knowledge extraction from published pathway figures is relatively rare and incomplete ^20;21;22^. In a pilot study of 4,000 pathway figures ^1^, we developed a custom OCR pipeline and identified over twice as many unique human genes as detected in the text by PubTator 23. The gene content extracted from this limited sample of pathway figures reproduced two-thirds of the database content at WikiPathways and included over a thousand human genes not previously annotated in pathway models. Remarkably, no two pathways were identical among this set of 4,000 figures. A wealth of novel and diverse pathway knowledge is essentially trapped in published pathway figures.

The goal of this work was to identify the human gene content in a comprehensive collection of published pathway figures, to characterize its biological relevance, and to increase meaningful, FAIR ^24^ access to this pathway knowledge resource. In the end, 65k pathway figures were found in publications from the past ^25^ years with over a million mentions of human genes identified in total. Of the 13.5k unique human genes identified, over a quarter had yet to be annotated in WikiPathways or Reactome databases. The biological relevance of the identified gene sets was assessed by performing enrichment analysis against annotated gene sets using Gene Ontology and an extensive disease ontology showing both diversity and depth. Finally, a series of usage examples demonstrate the potential of this content to enhance literature searches, elucidate the history of scientific discovery and support enrichment analyses.

## Results

We identified and characterized 64,643 pathway figures published between the years 1995 and 2019. Starting with 235,081 figures from a PMC image query that specified the 25-year date range and keywords covering diverse types of pathways, machine learning was applied to more precisely distinguish figures containing molecular interaction diagrams from those depicting other types of pathways (e.g., neuronal pathways) or pathway-related content (e.g., pathway enrichment results).

Relying solely on the linearly ranked PMC results (i.e., without subsequent machine learning steps) would have resulted a relatively *diluted* set of figures containing a high proportion of non-pathways. Two rounds of machine learning effectively *concentrated* actual pathway figures. The second and final round relied on a set of 15,406 figures manually classified by a domain expert to train a model distinguishing pathway figures from other figures with 91.88% precision, 91.88% recall, and a Matthews Correlation Coefficient 25 of 0.82. The resulting set of 64,643 pathway figures is defined as our “65k set” used in this study. The 65k set of pathway figures was ultimately assessed to consist of 94% pathways (±3% at 97% confidence) by manual classifying a random sample of 300 figures.

### Papers Containing Pathway Figures

Prior to any gene detection by OCR, the papers containing the 65k set of pathway figures were characterized by publicly available annotations. The pathway figures came from 56,095 papers authored by 216,542 unique authors and are published in 3,453 journals. Obviously, not all co-authors are involved in preparing a pathway figure in a given paper, but for comparison, the most successful effort to crowdsource pathway knowledge by the WikiPathways database has fewer than 800 unique authors.

The papers containing pathway figures can be characterized by paper-level annotations, for example disease ontology terms from Europe PMC and genes recognized in the text by PubTator ^23^. A subset of 29,187 (52%) pathway-figure containing papers had at least one disease ontology terms annotation from Europe PMC. The top 10 most frequent disease ontology terms annotating these papers are Cancer (39% of 29,187 papers), Infection (19%), Defects (15%), Tumor (9.3%), Diabetes (4.3%), Hypoxia (2.0%), Depression (1.3%), Obesity (0.8%), Ischemia (0.7%), Atherosclerosis (0.4%), and Other (9.2%). According to PubTator, 30,036 (53.5%) of pathway-figure containing papers had at least one gene found in the text (i.e., abstract and main body) with an average of 3.4 genes per paper. The top 10 genes found in the text of these papers are AKT1 (5.4% of 30,036 papers), MTOR (4.1%), TP53 (3.7%), MAPK1 (3.5%), TGFB1 (2.8%), PIK3CD (2.8%), EGFR (2.6%), TNF (2.4%), CTNNB1 (1.9%) and MAPK3 (1.7%). The majority of these genes match the dominant disease annotation for cancer-related biological processes.

### Genes in Pathway Figures

Having identified and characterized a set of papers containing pathway figures, the primary goal of this study was to extract their human gene content by an OCR pipeline customized for pathway figures 1. The approach targeted the most common ways authors refer to genes, proteins, complexes and families in pathway figures, leveraging current, alias and previous HGNC symbols as well as a curated collection of conventional bioentity names that have been mapped to official HGNC symbols. There is a wealth of information visually conveyed by pathway figures, including localization, reactions, cascades, cycles, co-factors, metabolites, drugs and other biological concepts. The most tractable and widely used content—even from fully-annotated pathway mod-els—is the gene content. The other components and concepts in pathway figures remain worthwhile pursuing in future work, but knowing the gene content on its own transforms a collection of static images into a resource with diverse research applications.

Of the 64,643 pathway figures identified by image classification, 58,962 (91%) figures had at least one human gene recognized by our pathway figure OCR pipeline. A total of 1,112,551 instances of human genes were recognized, consisting of 13,464 unique human NCBI Genes. On average, there were 18.9 genes recognized per figure, compared to only 3.4 genes recognized in the text of the same papers by PubTator. PubTator found a tenth as many genes (101,617) in the text of these same papers overall; only half of the papers (53.5%) mentioning one or more genes in the text. In our pathway figure OCR results, there were over 600 figures with more than 100 genes each. While many of the largest figures are interaction networks, the largest with 385 recognized genes is an augmented KEGG pathway where the authors properly listed the individual gene family and paralog members referenced generically in the original ^26^. At the other end, there was a long tail of just over 20k (37%) figures that had fewer than seven recognized genes.

The top 10 human genes identified in pathway figures representing unique families are MAPK1 (15% of 58,962 figures), AKT1 (14%), PIK3CA (10%), NFKB1 (8.9%), KRAS (7.6%), MTOR (7.5%), MAP2K1 (6.2%), TNF (5.6%), RAF1 (5.3%), and TP53 (5.1%). Compared to genes extracted from the text of the same set of papers containing pathway figures, the same trend of cancer-related biology dominates the content. In more detail, this top 10 set includes four of the top five genes found in the text of the same papers by PubTator: MAPK1, AKT1, MTOR and TP53. **Figure 1** presents the results of a set analysis performed on these overlapping genes found in text and figures, showing approximately 50% overlap with respect to text occurrences (i.e., half of the occurrences in text were also found in the pathway figures of the same papers) and overall a far greater number of occurrences in figures; the occurrence of combinations of these genes being almost exclusive to figures. Of the 13,464 unique human genes identified in the figures, half (6,564 or 49%) were not identified in the text of any of the papers by PubTator. Compared to pathway databases, over a quarter of the unique genes recognized in pathway figures (3,710 or 28%) were not present in either WikiPathways nor Reactome collections (as of January 2020). Clearly, pathway figures represent biological models that are not fully described in the text nor captured in curated pathway databases.

**Figure 1.**
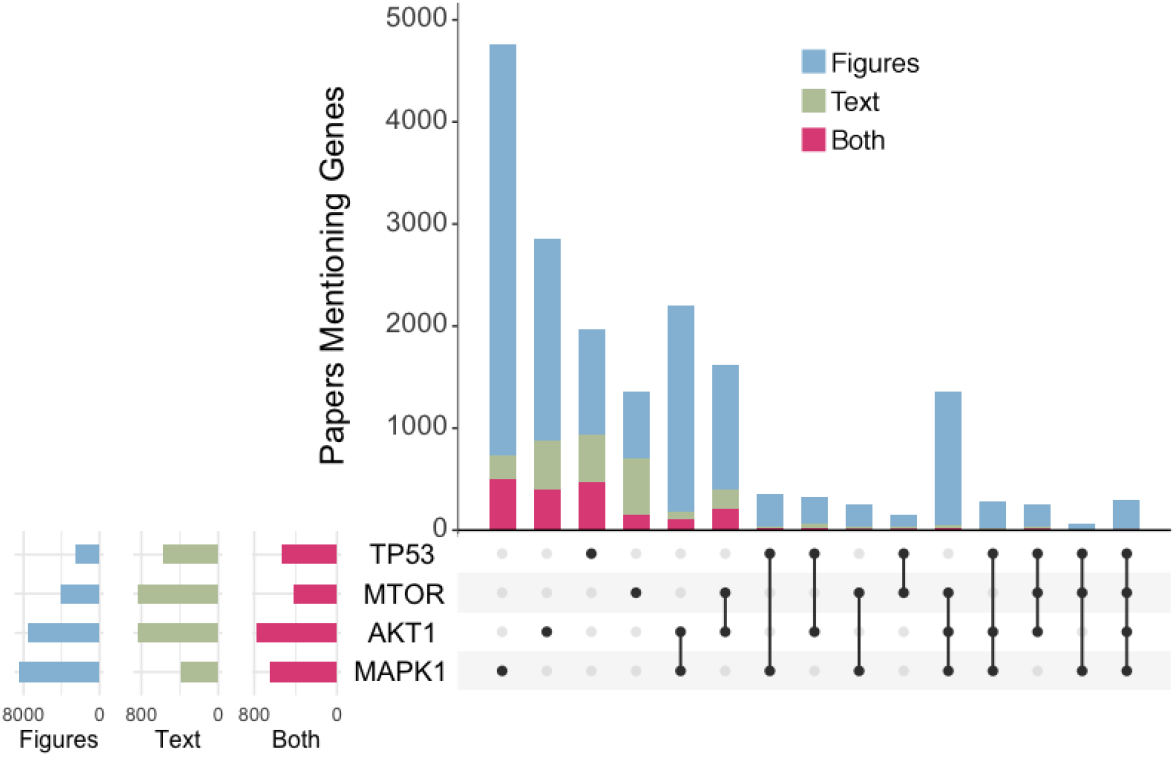
UpSet plot of the most common genes found in the text of pathway figure-containing papers. The stacked bar plot (top) shows the number of papers that include one or more of these genes in either a pathway figure (blue), the text via PubTator (green) or both (red). The matrix below the bar plot indicates which sets of genes are represented by each bar. The bar plots on the left show the total number of genes in either pathway figures, text or both using independent scales.

Concerned that the one-to-many mapping introduced by using bioentities might inflate unique gene counts, an “unexpanded” count was also calculated. Overall, there were 13,377 unique “unexpanded” NCBI Genes recognized, meaning that just 87 (0.6%) genes were *only* identified via bioentity expansion. Unexpanded gene counts were used to define subsets of figures for annotation and gene set enrichment analysis.

### Sets of Genes in Pathway Figures

Seeking to optimize across coverage, performance and interpretability, we defined a subset of figures with at least seven unique “unexpanded” NCBI Genes. Compared to the overall set, these 28,836 pathway figures contained 13,216 (98%) unique genes, thus retaining the coverage and novelty of the collection. Among these 28,836 pathway figures, 28,520 (99%) were significantly associated with at least one Gene Ontology (Biological Process) term by enrichment analysis, indicating general biological relevance. Using disease ontology annotations, we found 20,227 (70%) pathway figures significantly associated with at least one disease term, and 98% of disease terms in the ontology (157/160) were represented by one or more figures. The top 10 disease terms associated with these pathway figures were Cancer (42% of 20,227 figures), Juvenile rheumatoid arthritis (7.8%), Ovarian cancer (6.8%), Primary hyperaldosteronism (5.0%), Aortic aneurysm (3.5%), Alopecia areata (3.1%) Primary cutaneous amyloidosis (2.7%) Melanoma (2.3%) Alzheimer’s disease (2.1%) Rheumatoid arthritis (2.1%), and Other (23%) (**Figure 2.B**). Compared to the disease annotations on the parent papers, both lists are clearly dominated by Cancer and then spread to cover a wide range of diseases. The gene information provided by the pathway figures, however, allows for enrichment against higher resolution disease ontologies, as well as to any gene set-based ontology or resource, such as Gene Ontology, OMIM, MSigDB or even other pathway databases like WikiPathways and Reactome.

**Figure 2.**
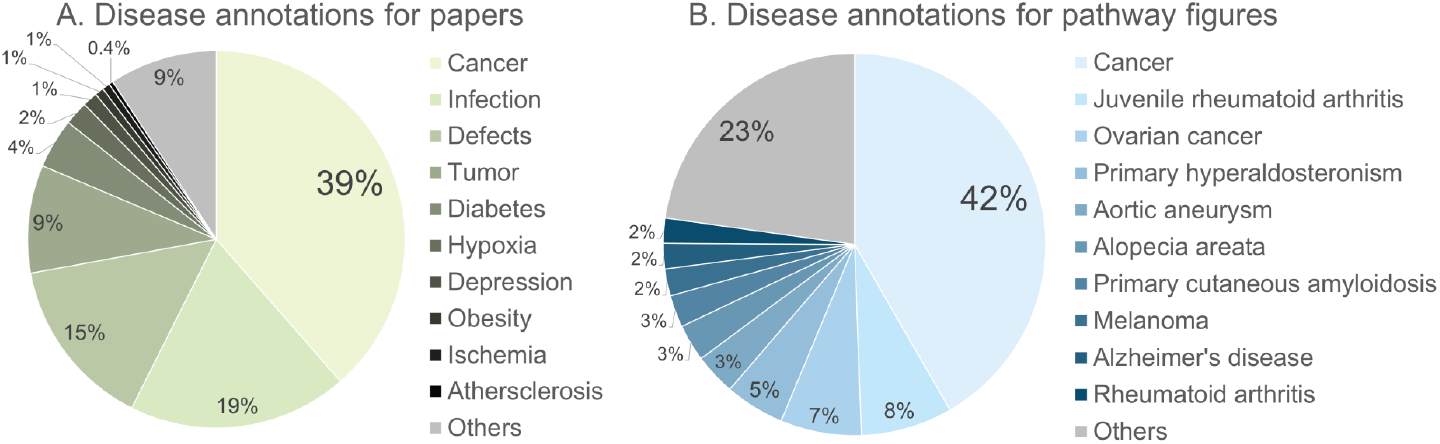
Disease annotations for pathway figures and their parent papers. A. Top 10 disease annotations from Europe PMC for 29,187 of the papers containing pathway figures. B. Top 10 enriched disease annotations for 20,227 pathway figures based on extracted gene content.

### Utility of Pathway Figures

The initial characterization of published pathway figures and their gene content revealed a novel resource of relevant pathway knowledge that is practically inaccessible to researchers. The following examples demonstrate how this resource could be utilized in a variety of applications.

### Searching Scientific Literature

With over 600,000 papers added to PMC in the last year, researchers are resigned to merely sampling the work most relevant to them via search engines, feeds, subscriptions and recommendations. While figure captions are accessible to text-based processing and indexing, the actual contents of figures remain hidden to any commonly available search engine (e.g., PubMed, PMC, Europe PMC or even Google). The systematic identification of genes in pathway figures enables access to this content through new and existing tools.

#### Literature search tools

Search engines commonly index papers by the genes found in the abstract, body and caption text. Europe PMC goes further by supporting community-contributed mappings between genes and papers (https://europepmc.org/annotations). While these typically derive from text-based processing, the same deposition and integration system could be used to accept gene-paper mappings based on figures. Querying one or more genes at Europe PMC would then return paper results containing both text and figure references to those genes.

As another example, the Chan Zuckerberg Initiative is working on a new literature feed service called Meta (https://meta.org; in open beta) that processes the latest publications and preprints on a daily basis. The gene-paper mappings from pathway figures would be a natural fit for an indexing system that links papers via their contents. Furthermore, the characterization of gene sets as demonstrated above could provide mappings from papers to disease ontology terms and other gene-based and pathway-based annotations. A feed service that could produce a regularly updated set of papers that contained relevant pathway figures would be a welcome innovation to researchers attempting to stay abreast of scientific literature.

#### Interactive pathway figure app

We produced an online tool using R Shiny (https://gladstone-bioinformatics.shinyapps.io/shiny-25years) to enable filtering, searching and viewing the full collection of 65k pathway figures by enriched disease terms, genes, date and various publication metadata fields (**Figure 3**). Organized into three stages, the first stage offers auto-complete fields to define OR-based filters for disease annotations, gene content and publication years, and displays bar plots of the top 40 disease ontology terms, top 40 human genes and publication dates represented by the currently filtered set of figures. The second stage displays a paginated table view of the currently filtered set of figures, each row representing a pathway figure and its parent paper. The columns can be used to sort and query within the table to further refine the current set. Selecting a row in the table will update the third stage, which displays the pathway figure, a link to PMC and table recognized genes. The gene table includes the symbol found in the figure and the source of the lexicon it matched along with the official HGNC symbol and NCBI Gene identifier. This tool provides a way to easily query figures of interest given a set of genes or topic. For example, in the preparation of this manuscript the tool was used for the section on History of Scientific Discovery relating to the Hippo Signaling Pathway.

**Figure 3.**
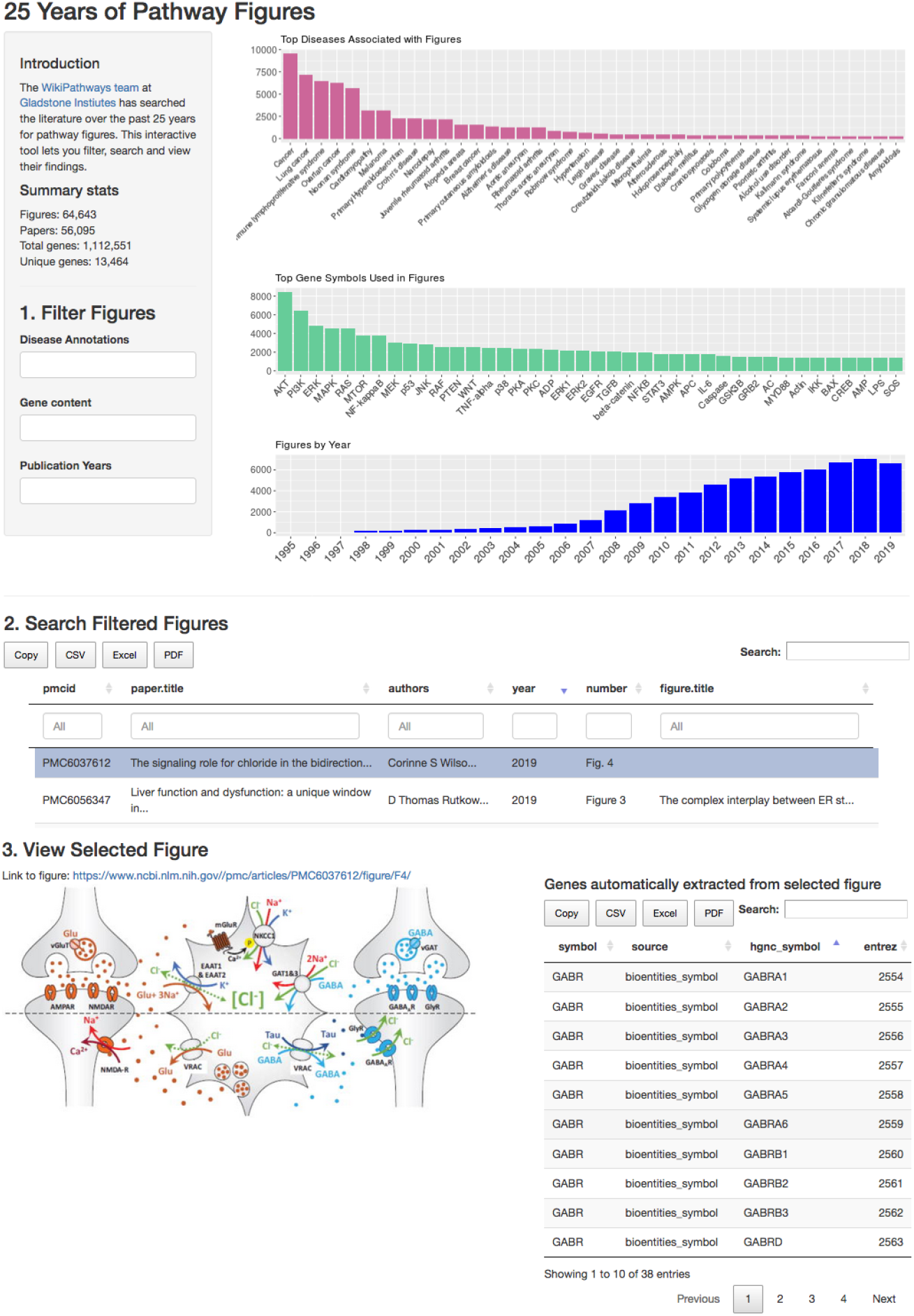
Screenshot of interactive query tool for pathway figures. The tool allows for easy querying of the full set of 65k pathway figures by any combination of gene content, publication year, disease annotations, figure and paper title, authors etc.

As a topical demonstration of the tool, a second version was produced focusing on COVID-19 related pathways as defined by the COVID-19 Open Research Dataset (]https://gladstone-bioinformatics.shinyapps.io/shiny-covidpathways). There are 221 pathway figures in this collection that can be rapidly queried and viewed by the same three-stage procedure described above and in **Figure 3**. This tool has already proven useful in building SARS-CoV-2 pathways at WikiPathways (http://covid.wikipathways.org) as part of the COVID-19 Disease Map initiative (https://covid.pages.uni.lu/map_curation) ^27^.

#### Knowledge graph query paths

This source of annotated pathway information is also finding utility in an advanced platform for distributed knowledge integration. The BioThings Explorer platform includes a collection of APIs semantically defining inputs and outputs that comprise a knowledge graph (https://biothings-explorer.readthedocs.io) ^6^. The platform also includes an engine that supports queries that traverse paths through the graph, e.g., *drugs* that bind *components of pathways* associated with a *disease of interest*. By defining an API that recognizes standard paper and gene identifiers in a JSON file export of pathway figure-based gene sets, the 1.1M gene-paper links from this study can be used to bridge query paths to other content in the knowledge graph. The addition of disease annotations and future extraction of metabolites, drugs and concepts from the OCR results will establish additional bridges and reinforce paths inferred by computational reasoning. This work is in active development as part of the NCATS Biomedical Data Translator program, targeting critical use cases, such as drug repurposing (https://ncats.nih.gov/tidbit/tidbit_04.html).

### History of Scientific Discovery

Given the 25-year span of this new pathway resource, it was natural to reflect on the role of pathway figures in scientific discovery. The R Shiny app previously described can be used by any researcher or historian to investigate the story of particular diseases and genes from a pathway perspective. Tracing the development of the Hippo signaling pathway as one example demonstrates this potential (**Figure 4**).

**Figure 4.**
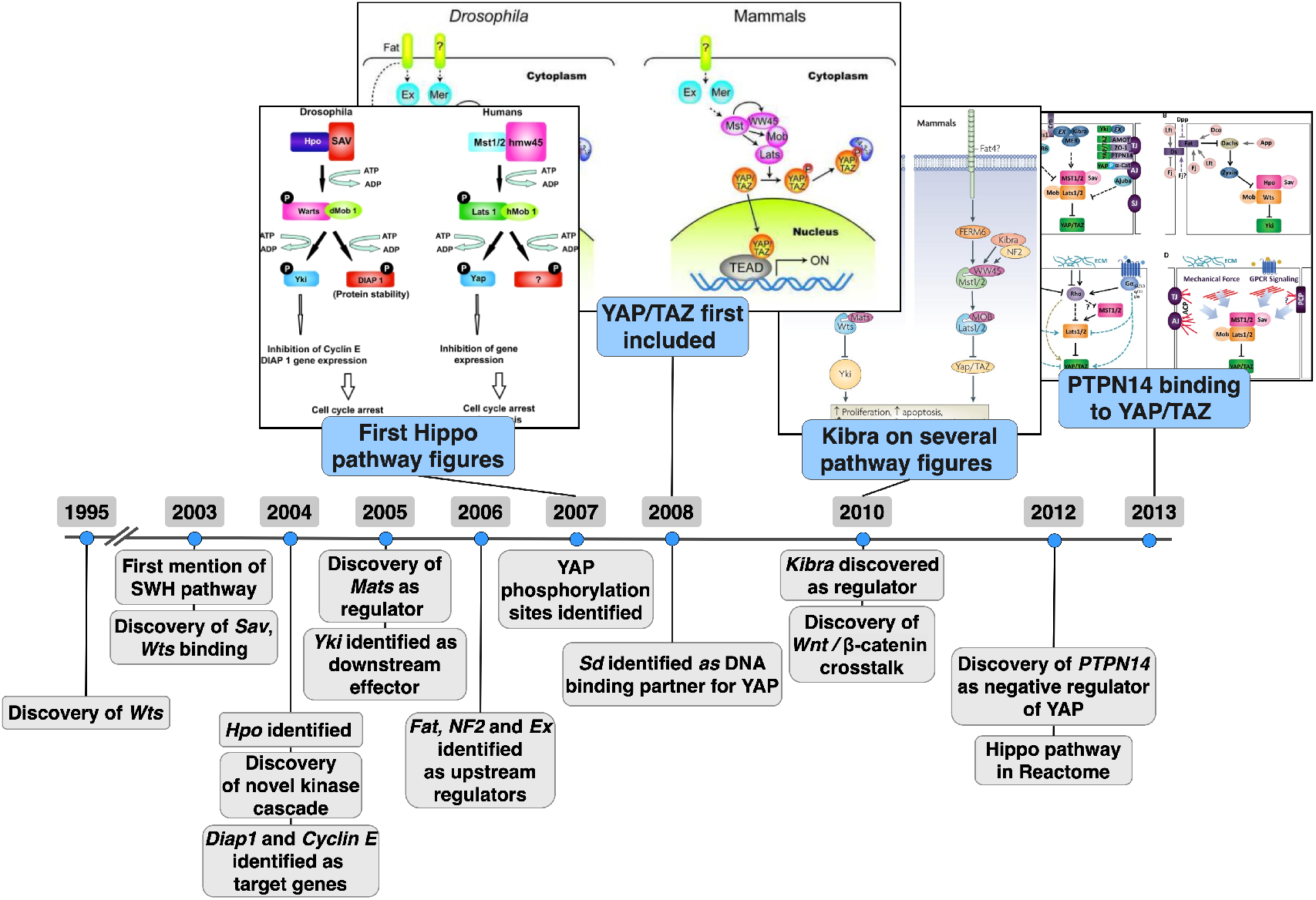
Major milestones in the development of the Hippo signaling pathway. A timeline spanning 1995 to 2013 focuses on the discovery of the components and processes of the Hippo signaling pathway (below timeline), illustrated by representative pathway figures that grow in complexity and detail (above timeline). The events depicted in grey below the timeline were collected mostly from Kim *et al.* ^28^. Drosophila and human orthologs are Wts:LATS1/2, Sav:SAV1, Hpo:MST1/MST2(STK3), Mer:NF2, Fat: FAT1, Ex: FRMD6, Yorkie/Yki:YAP(YAP1)/TAZ(WWTR1), Sd:TEAD1-4, Kibra:WWC1.

The Hippo signaling pathway controls organ size in animals by regulation of cell proliferation and apoptosis. The components of the Hippo signaling pathway are highly conserved ^29^, and many of the early discoveries were made via Drosophila genetic screens. The Hippo signaling pathway includes a central kinase signaling cascade, where MST1/2 (Hpo) phosphorylates LATS1/2 (Wts), which activates it. Activated LATS1/2 phosphorylates YAP/TAZ (Yorkie/Yki), leading to its inactivation and degradation in the cytoplasm. When activated, YAP/TAZ translocates to the nucleus and binds to several transcription factors, including TEADs, leading to transcription of proliferation and survival genes. Phosphorylation of LATS1/2 is facilitated by binding to SAV and MOB1. The Hippo pathway can be activated by many different stimuli including cell density and polarity, mechanical sensation and soluble factors via upstream regulators WWC1 (Kibra), NF2 etc. Cross-talk with multiple pathways is known, including TGF-*β*, Notch and Wnt signaling. The Wts gene (LATS1 in humans) was discovered in Drosophila in 1995^30;31^, and the first mention of a pathway involving Wts, Sav and Hpo was in 2003^32;33^, initially termed the Salvador/Warts pathway. From 2003 on, several discoveries were made which further defined the pathway components and process ^28^.

In our 65k pathway figure collection, the first published figures representing the Hippo signaling pathway appeared in 2007^34;35^, more than a decade after the initial discovery of the central Wts gene, and after the core components were described in the literature (**Figure 4**). The early published pathway figures are sparse and some even include question marks for components that are not yet known ^34^. Discoveries of specific components are in some cases followed by a pathway figure from an independent publication, adding that component to the pathway. For example, discovery of the binding of PTPN14 to YAP in 2012^36^ is followed in 2013 by a pathway figure showing the interaction ^37^. Another interesting observation is related to the Kibra gene (WWC1 in humans), which was first characterized in a yeast two hybrid screen in 2003^38^. It was investigated in a variety of contexts (cytoskeleton, memory function etc), and in 2010 Kibra was shown to be an upstream regulator of the Hippo pathway ^39^. Interestingly, there were no pathway figure hits for Kibra before 2010, but from 2010 on the number of pathway figures including Kibra has grown steadily, with the vast majority of them representing the Hippo pathway. Yorkie (YAP/TAZ in humans) was first indicated as the transcriptional activator of the Hippo pathway in 2005^40^ and it was subsequently found that phosphorylation of YAP at S127 inhibits transcriptional activity by retaining YAP in the cytoplasm ^41^. These critical findings were followed in 2008 by the first pathway figure showing the details of YAP/TAZ signal transduction.

The first representation of the Hippo signaling pathway in a major pathway database was in 2012, when it was added to Reactome (https://reactome.org/content/detail/R-HSA-2028269). In our results, there were 31 representations of the Hippo pathway in published literature spanning the 13 years prior to its database entry in 2012 (1999-2011), indicating that extracting pathway knowledge from published figures is important in capturing emerging pathway knowledge.

Trends for all pathways and their gene content can also be explored. For example, by using data gathered from OMIM ^42^, the time spanning the initial cloning of a gene and its first appearance on a pathway can be determined. A decade in the case of the Wts (LATS1/2 in human) gene and the Hippo signaling pathway, the cloning-to-pathway time spans for all 13,464 genes has an overall median of 12 years. By comparison, the time spanning the initial cloning of a gene and its first biochemical feature characterization also has a median of 12 years.

### Pathway Figure Enrichment Analysis

The 65k set contains more unique human genes and greater contextual depth than any pathway database, providing an interesting new resource for pathway analysis. However, to make the set usable for enrichment analysis like overrepresentation and gene set enrichment analysis, there are aspects of gene content and redundancy to consider.

Even though certain methods normalize enrichment scores for gene set size, the process is not accurate for extremely small or extremely large gene sets. While large sets are not an issue in our pathway figure-based gene sets, approximately half of the figures have fewer than 10 genes. Filtering the 65k set with a cutoff of at least 10 unique genes, a set of 32,277 (49%) pathway figures can be defined for use in enrichment analysis. By contrast, only 1,072 (1.9%) papers had 10 or more genes identified in the text by PubTator. Also of note, this set of 49% of the largest pathway figures retains 97% (13,153) of the unique human genes found in the overall 65k collection. Among the set of 32,277 pathway figures identified for use in enrichment analysis, there were 878 figures that shared the exact same gene content with at least one other figure and 4,937 figures entirely contained by one or more other pathway figures.

In order to assess the redundancy and hierarchical structure among pathway figures in more detail, a sample subset of 55 Hippo Signaling pathway figures was clustered by gene overlaps (**Figure 5**). The overlap (intersection / number of genes in gene set) and Jaccard index (intersection / union) ^43^ were calculated between each pair of gene sets. No two pathways in this set were identical, but many pathway figures contained the contents of smaller pathway figures, analogous to the nesting of Biological Process terms in Gene Ontology. The “Core” cluster (red), for example, contains four small figures each with the defining set of Hippo Signaling pathway genes, which are an essential component of all the figures in this set; thus, the high scores across top four rows. The “Meta” cluster (purple) has ^25^ large figures associating multiple pathways with Hippo Signaling. Though there is much lower similarity for this cluster, a few bright subclusters along the diagonal indicate sets of pathway figures with high mutual overlap. The “Plus” cluster (green) has ^26^ small-to-medium figures that contain just a few additional genes interacting with the core Hippo Signaling pathway. In the context of an enrichment analysis, any of these pathways have the potential to provide a highly specific result with greater context and interpretability than just a single, so-called canonical, Hippo Signaling pathway. This potential argues against additional pre-filtering of the pathway figure set. Furthermore, many enrichment tools already employ post-filtering of results by, for example, a Jaccard distance measure to handle ontologies with even greater redundancy and higher degree of nesting. The same approach could optionally be applied to enrichment results from these pathway figure-based gene sets as part of a researcher’s exploratory analysis.

**Figure 5.**
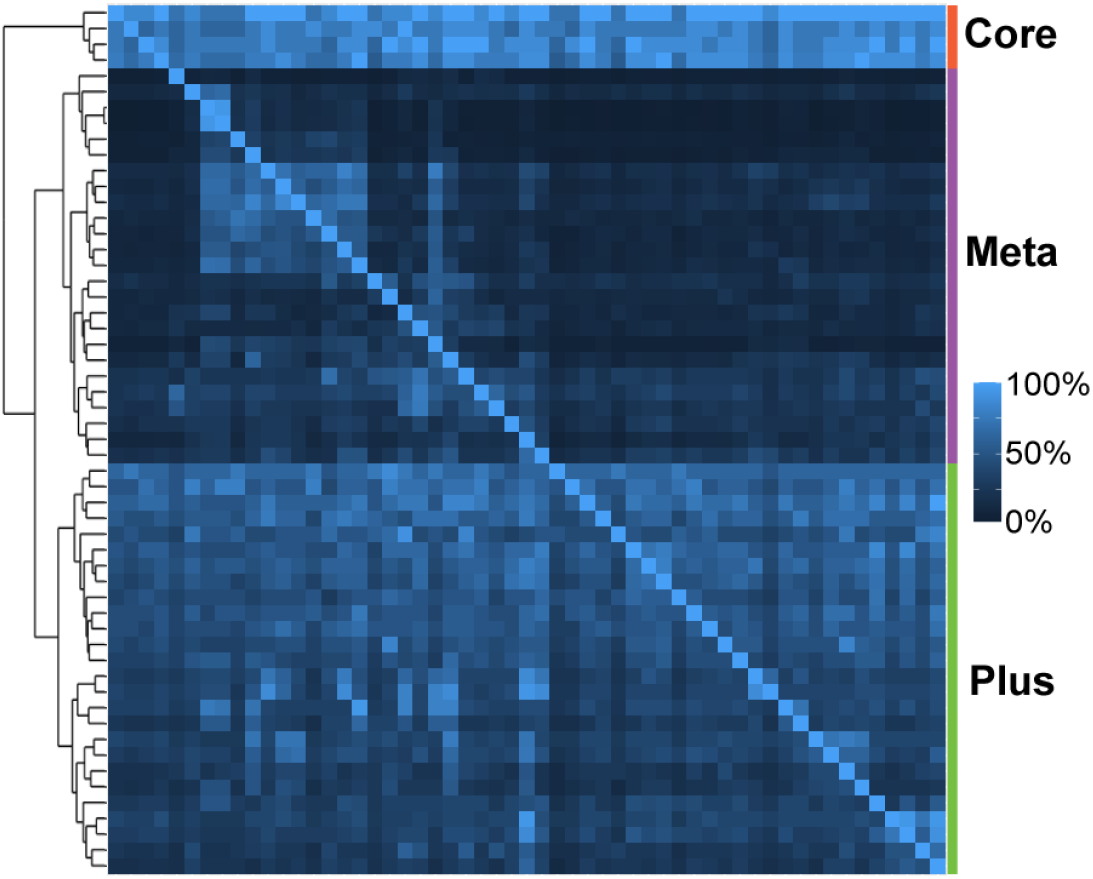
Overlap among Hippo Signaling pathway figures. A matrix view of the directional overlap among 55 pathway figures containing at least five core genes involved in hippo signaling. The dendrogram (left) identifies various levels of clustering. The top three clusters are annotated by labels and a color bar for reference (right).

## Discussion

There is a vast resource of pathway knowledge trapped in the form of published pathway figures. We have identified these figures and extracted their gene content to make this resource accessible and to demonstrate its potential to enhance biomedical research. We found 64,643 pathway figures in publications dating from 1995 to 2019 and identified 1,112,551 occurrences of human genes in total (13,464 unique NCBI Genes). Much of this content represents novel pathway knowledge that is present neither in the text of papers, nor in pathway databases. The extraction of drugs, metabolites and disease terms from these same pathway figures is now a more tractable project, as is the identification of genes from other species, by expanding the lexicon used to match against the OCR results.

Despite unlocking the contents of published pathway figures, this work only partially mitigates gross deficiencies in current pathway knowledge representation and communication. *Oh, the figures we have seen!* Having scanned many thousands of pathway figures as a collection, the most obvious point to make is that standardization is desperately needed. Standards for pathway models have been around for decades ^44;45;46^ and have been implemented in a variety of freely available software tools ^47;48;49;50;51^. Likewise, standard practices for the deposition and sharing of scientific models are well established (e.g., sequences, structures and ontologies). We recommend that authors make use of these standards and we implore reviewers, editors, journals and funders to encourage and enforce the application of good scientific publishing practices to pathway knowledge. This recommendation applies generally to the publication of pathways, networks and other models of system biology consisting of identifiable entities and their relationships. By using proper modeling tools, pathway knowledge can be databased, indexed, shared and used more effectively and FAIRly ^24^.

In the meantime, the *post hoc* extraction of knowledge from published pathway figures can serve to make this content more findable, accessible, interoperable, and reusable. The gene sets extracted from these figures can be indexed to enhance literature searches, they can create and reinforce links in knowledge graphs, they can inform the historiography of gene and pathway discovery, and they can enable pathway figure-based enrichment analysis. This work also enables the prioritization of pathway figures (e.g., by novel gene content) to be curated as proper models in pathway databases. The interactive pathway figure tool allows anyone to explore the complete set of 65k figures by various metrics and metadata (https://gladstone-bioinformatics.shinyapps.io/shiny-25years).

## Methods

### Collection of Figures

A PMC image query URL was composed, specifying a date range starting at 1995-01-01. Exploratory queries previously identified 1995 as the first year with accurately indexed pathway figures. Keywords in the query were specified as an OR set of the following pathway types together with the word *pathway*: *signaling, signalling, regulatory, disease, drug, metabolic, biosynthetic, synthesis, cancer, response*, and *cycle*. These were determined based on exploratory queries and referencing the first two levels of pathway ontology terms ^52^. The query returned just over 235,000 figure images, which were retrieved by an HTML-scraping script, along with metadata pertaining to the figure and parent paper: PMCID, paper title, paper citation, publication year, figure filename, figure URL, figure number, figure title and figure caption. The collection was filtered for unique entries and publication dates spanning the 25-year period of 1995-01-01 to 2019-12-31. It is worth noting that while the query was performed on 2020-01-31, the results for 2019 are not expected to be complete since many journals unfortunately wait six months to a year to make their content openly accessible.

#### Limitations

Web scraping the results of a PMC image query is inefficient and imprecise. It is likely that many pathway figures are missing from the results due to incomplete keyword listing and database indexing. At the same time, the query results included many non-pathway figures. Given the ranked order of images provided by provided by PMC, we manually checked the percentage of *actual* pathway figures at three points: the first thousand figures (of 235k) contained 66.3% pathways, the middle thousand contained 35% and the final thousand contained fewer than 10% (**Figure 6.A**). The order of query results from PMC was thus informative, but not sufficient to distinguish pathway and non-pathway figures.

**Figure 6.**
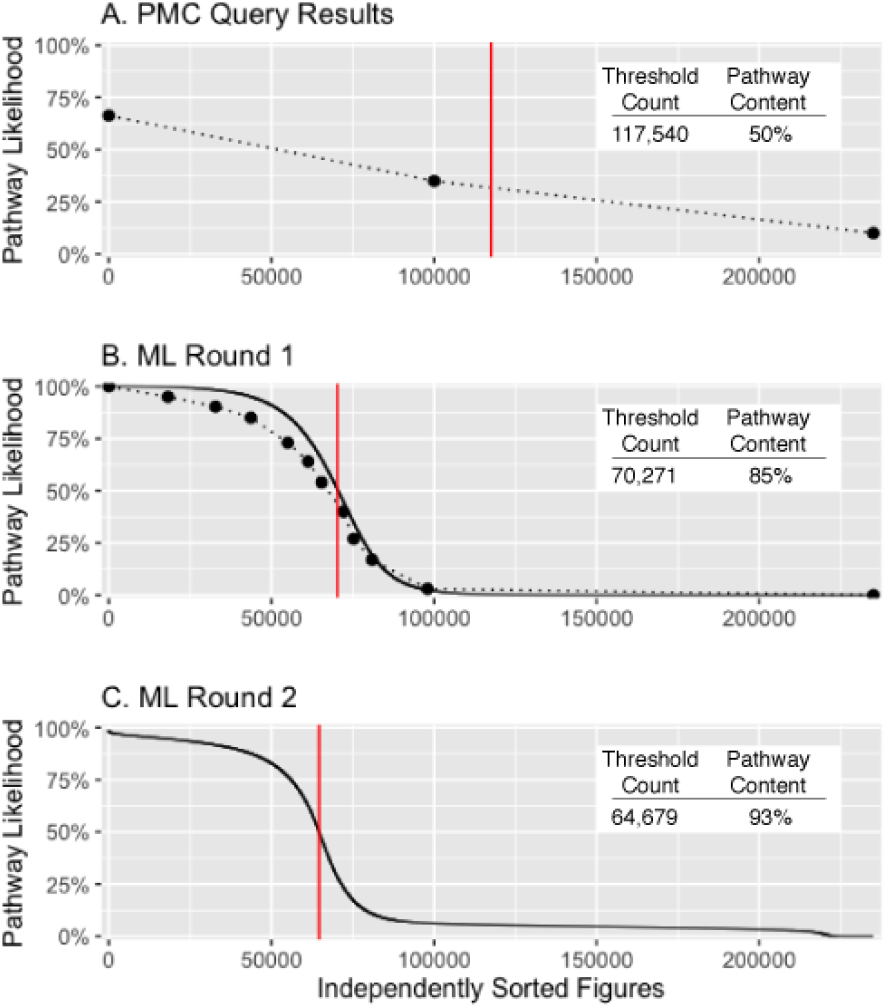
Machine learning performance. Classification of pathway figures assessed in (A) original PMC-sorted query results and (B) first and (C) second rounds of machine learning results independently sorted by estimated pathway likelihood (y-axis). Dots in A and B (connected by dotted lines) indicate where figures were sampled along their sorted indices for manual classification. These data were used in training subsequent rounds of ML. The solid black curves in B and C indicate the prediction of the learned model from each round of ML. The red vertical lines indicate the threshold used to define a set of pathway figures at each stage and assess their count and actual pathway content (inset tables). The final set of 64,643 pathway figures replaced 1.2% of known false positives and false negatives based on manual classification and was estimated to contain 94% pathway content.

### Classification of Figures

To properly identify pathway figures among the PMC image query results, two rounds of machine learning were performed using Google Cloud AutoML Vision. The AutoML service was accessed and controlled via a REST API and a web-based dashboard. The first model was trained on 2,000 manually classified figures selected from high, middle and low relevance ranges based on PMC query result ranking (**Figure 6.A**, dots). The service randomly split the provided figures into three sets: 80% training, 10% testing, and 10% validation. The manual classification was performed by a domain expert relying on their own organic neural network (a.k.a. brain) trained on 15 years of experience creating, curating and using pathway diagrams in biomedical research. For the purposes of this OCR-based project, figures were considered pathways if they described a biological process or set of interactions involving identifiable genes and proteins. Molecular interaction networks and developmental processes were thus included, while cellular diagrams that did not name genes or proteins were excluded. The AutoML service evaluated the performance of the first model at 88.42% precision and 91.3% recall at a 50% confidence threshold. The model was then applied to the complete set of 235k figures to obtain pathway likelihood scores (**Figure 6.B**, solid line). Additional figures were sampled along the full distribution of scores and manually classified (**Figure 6.B**, dots). The total actual pathway content above the threshold (red line) was estimated to be 85%. Given these results, a second model was trained on combined set of 15,406 manually classified figures and assessed to perform at 91.88% precision and 91.88% recall at a 50% confidence threshold, resulting in a steeper transition (i.e., fewer uncertain calls) and increased accuracy at the extremes (**Figure 6.C**). A Matthews Correlation Coefficient of 0.82 was calculated 25. 64,679 figures had a predicted pathway likelihood of 50% or greater from the second round of machine learning (red line). Finally, 383 false negative pathways were added back and 419 false positive non-pathways were removed based on prior manual classification, resulting in the set of 64,643 pathway figures used in this study.

#### Limitations

A random sample of 300 figures from the final set was manually classified to estimate the proportion of actual pathway figures at 94% (±3% at 97% confidence). Of the 18 figures classified as non-pathways, two were composite figures that included a pathway as a minor panel in figure that contained a lot of non-pathway content. Composite figures were typically excluded from the “pathway” training sets in our manual classification in order to avoid recognizing gene occurrences in figure elements unrelated to pathways, so these were conservatively included in the false positive count. Of the remaining 16 non-pathways, only three had three or more genes subsequently detected by our pathway figure OCR pipeline (see next section), suggesting the majority of false positives could be effectively ignored.

### Identification of Genes in Pathway Figures

The set of pathway figures was then fed into our pathway figure OCR pipeline 1. Briefly summarized here, the pipeline’s main components are word isolation, transformation and lexicon matching. A series of custom transforms were applied to newline-and space-delimited words provided by the OCR output. Transforms included normalization of characters, substitutions and expansions. After each round of transformation a match is attempted against a lexicon of human genes including HGNC symbols (official, aliases and previous) and bioentities (https://github.com/wikipathways/bioentities). The pipeline output is a table of pathway figures associated with sets of recognized genes with human NCBI Gene identifiers. Paper and figure metadata including titles and captions are maintained along with additional information such as the raw OCR results, the normalized and transformed symbols and the lexicon source that was matched.

#### Limitations

There were three non-composite false positive cases that contained three or more human genes: a diagram of nerve fibers with gene markers ^53^, a meta-network of gene-named pathways (e.g., IL-6 signaling) ^54^, and a figure containing three-letter amino acid codes (e.g., Tyr, His, Met) that happen to match gene aliases ^55^. The first two cases are not egregious in that relevant biology is still being detected; it is just not in the context of a pathway diagram as defined here. The last case is the only one that is a problematic false positive (i.e., mistakenly identifying an amino acid as a gene). The pathway figure OCR pipeline was also limited by the lexicon of human genes. Many of the pathway figures with zero or small numbers of recognized genes were for other species, e.g., Drosophila signaling pathways, yeast networks and microbial metabolism. The lexicon can be expanded in the future to include other species, other types of molecules and other biological concepts.

### Characterization of Pathway Figures

Annotations on the 56,095 papers containing pathway figures were retrieved from the PMC query site and web services, including authors, journal titles, paper titles, paper identifiers, publication dates, figure titles, hyperlinks and captions.

Disease annotations for 29,187 papers were available from Europe PMC and collected using the *europepmc* R package ^56^. A less redundant list of top 10 terms was made by excluding previously counted papers and re-sorting by disease term frequency. Singular and plural forms of the same disease term were combined, e.g., “Tumor” and “Tumors”. After identifying the top ten, remaining publications were counted as “other”.

Gene associations for 30,036 papers were available from PubTator and downloaded as NCBI Gene-to-PMID mappings from the PubTator FTP Service. A PMID-to-PMCID mapping file from the PMC FTP Service enabled comparison with our PMCID-indexed pathway figures and extracted gene sets.

Gene Ontology and disease annotations for pathway figures were determined by performing enrichment analysis on the sets of figure-extracted genes against ontology-associated gene sets. The source of disease annotations was the “knowledge” channel of the DISEASES database ^57^ filtered for disease terms with seven or more associated genes, resulting in a set of 160 disease terms in total with 5,088 gene associations. Top 10 disease term lists were made less redundant by excluding previously counted figures (i.e., figures associated with more than one disease term) and re-sorting by disease term frequency.

## Data Availability

- OCR pipeline code repository:
https://github.com/wikipathways/pathway-figure-ocr
- Interactive pathway figure search tool:
https://gladstone-bioinformatics.shinyapps.io/shiny-25years

## Author Contributions

KH, AR and ARP conceived of the project. AR implemented the majority of the AutoML and OCR pipelines. All authors conducted analyses presented in the paper. All authors contributed to writing and editing of the manuscript.

